# Aneuploidy-driven genome instability triggers resistance to chemotherapy

**DOI:** 10.1101/2020.09.25.313924

**Authors:** Marica Rosaria Ippolito, Valentino Martis, Christy Hong, René Wardenaar, Johanna Zerbib, Diana C.J. Spierings, Uri Ben-David, Floris Foijer, Stefano Santaguida

**Affiliations:** Department of Experimental Oncology at IEO, European Institute of Oncology IRCCS, Via Adamello 16, 20139 Milan, Italy; European Research Institute for the Biology of Ageing, University of Groningen, University Medical Center Groningen, 9713 AV, Groningen, the Netherlands; Department of Human Molecular Genetics and Biochemistry, Faculty of Medicine, Tel Aviv University, Tel Aviv, Israel; Department of Oncology and Hemato-Oncology, University of Milan, Via Santa Sofia 9/1, 20122 Milan, Italy

**Keywords:** aneuploidy, cancer, genome instability, chemotherapy, drug resistance

## Abstract

Mitotic errors lead to aneuploidy, a condition of karyotype imbalance, frequently found in cancer cells. Alterations in chromosome copy number induce a wide variety of cellular stresses, including genome instability. Here, we show that cancer cells might exploit aneuploidy-induced genome instability to survive under conditions of selective pressure, such as chemotherapy. Resistance to chemotherapeutic drugs was dictated by the acquisition of recurrent karyotypes, indicating that gene dosage, together with mutational burden, might play a role in driving chemoresistance. Thus, our study establishes a causal link between aneuploidy-driven genome instability and chemoresistance and might explain why some chemotherapies fail to succeed.

## Introduction

Chromosome mis-segregation leads to abnormal karyotypes, a condition known as aneuploidy (Santaguida and Amon 2015). The presence of aneuploid karyotypes affects several processes and have many cellular consequences, including genome instability, metabolic alterations and proteotoxic stress (Chunduri and Storchova 2018). In humans, aneuploidy is the primary cause of spontaneous abortions and leads to severe developmental defects, such as those present in Down syndrome patients (trisomy 21) (Roper and Reeves 2006). Importantly, the aneuploid state is highly prevalent in cancer and the presence of aneuploid karyotypes correlates with poor patient prognosis (BenDavid and Amon 2019) and resistance to chemotherapy (Gómez-Miragaya et al. 2019; Andor et al. 2015; Birkbak et al. 2011). Chemotherapy is a central therapeutic strategy for most cancer patients and resistance to therapeutic drugs leads to the failure of this treatment. Thus, studying and understanding chemoresistance is a major challenge in cancer biology. Because of this, there are several ongoing efforts concentrated on understanding the contribution of cell intrinsic factors - such as genetic alterations and epigenetic changes – as well as cell extrinsic stimuli - such as cytokines and growth factors – as major players responsible for drug resistance (Vasan et al. 2019). Recent papers identified an increased resistance of aneuploid cancer cell lines to multiple chemotherapies, and to drugs in general (Cohen-Sharir et al. 2020). However, very little is known about how and why aneuploidy and the ensuing genomic instability impacts the therapeutic outcome. An intriguing hypothesis is that aneuploidy and genome instability provide phenotypic variation, thus increasing heterogeneity within a tumor and driving the ability of cancer cells to adapt to stressful conditions, including chemotherapy. In agreement with this idea, evidence from experiments in yeast has shown that when cells are cultured under strong selective pressure, aneuploid karyotypes can arise as an adaptive mechanism of survival (Rancati et al. 2008; Selmecki et al. 2006). Likewise, in mammalian cells, it has been proposed that aneuploidy is able to provide a proliferative advantage under selective conditions (Rutledge 2016). Furthermore, copy-number intratumor heterogeneity (ITH) has been associated with worse overall survival in patients (Jamal-Hanjani et al. 2017; Andor et al. 2015). These observations indicate that aneuploidy may be specifically exploited by eukaryotic cells to thrive under unfavorable growth conditions and suggest that karyotypic heterogeneity might be a vital resource for cancer cells during therapeutic drug treatments.

To formally test and study the relationship between aneuploidy and chemoresistance, we elevated chromosome mis-segregation rate in a panel of cancer cells prior to exposing them to common clinical chemotherapeutic drugs. We found conditions in which induction of mitotic errors had beneficial effects in the presence of chemotherapeutic agents.

Importantly, single cell sequencing analysis revealed specific karyotype recurrence in resistant cells. We speculate that aneuploidy-induced genome instability might trigger therapeutic drug resistance through the expansion of karyotype heterogeneity and subsequent convergence onto specific, favorable karyotypes that are crucial for cell survival. Clonal karyotypic evolution ensures cell viability through changes in the dosage of specific gene products, such as the therapeutic target, drug efflux pumps or metabolic enzymes. Therefore, our results provide the first direct evidence for a role of aneuploidy in driving adaptability during chemotherapy. Finally, giving the fact that there are ongoing clinical trials involving agents that elevate chromosome mis-segregation rate (Pauer et al. 2004; Mason et al. 2017; Wang et al. 2019), our study strongly suggests that such a pharmacological approach might not be invariably detrimental for cancer cells, but could actually promote cancer cell survival in some cases, highlighting the need to identify the exact conditions in which patients might benefit from such drugs.

## Results and Discussion

To begin to investigate whether and how elevation of chromosome segregation errors and the resulting chromosomal instability (CIN) provides a proliferative advantage under conditions of selective pressure, we first induced chromosome mis-segregation in a panel of cancer cell lines by transiently treating them with reversine, a small-molecule inhibitor of the mitotic kinase Mps1 (Santaguida et al. 2010). We then removed the drug and monitored their proliferation over time either in the absence (Figure 1A-C) or in the presence of a chemotherapeutic agent (Figure 1D-G). In agreement with previous reports (Santaguida et al. 2015; Williams et al. 2008; Sheltzer et al. 2017; Stingele et al. 2012; Santaguida et al. 2017; Tang et al. 2011), induction of CIN led to decreased proliferation (Figure 1B, C). To test the effects of CIN on cell proliferation in the presence of a chemotherapeutic agent, cancer cell lines were exposed to a battery of chemotherapeutic drugs, after reversine removal (Figure 1D, E). We used a panel of cancer cell lines from different tissues of origin, including colon, lung, pancreas and skin and continuously exposed them for 6 weeks to anticancer agents routinely used in the clinic (Figure 1D, E). Among tested conditions (Figure 1E), we found combinations in which pretreatment with reversine provided an advantage, leading to cell survival and colony formation at the end of our experimental protocol (Figure 1E). A showcase of this behavior is given by the nonsmall cell lung cancer (NSCLC) NCI-H1975 treated with the topoisomerase I inhibitor Topotecan (Figure 1F). Under these conditions, induction of CIN through reversine pulse provided permissive conditions for cell survival in presence of Topotecan. This was not limited to the particular combination of NCI-H1975 with Topotecan, but we also found other examples, including the colorectal cancer cell line RKO and the pancreatic cancer cell line PANC1 treated with the thymidylate synthase inhibitor 5-Fluorouracil, the malignant melanoma cell line A375 treated with the B-Raf inhibitor Vemurafenib (Figure 1E, G).

**Figure 1.**
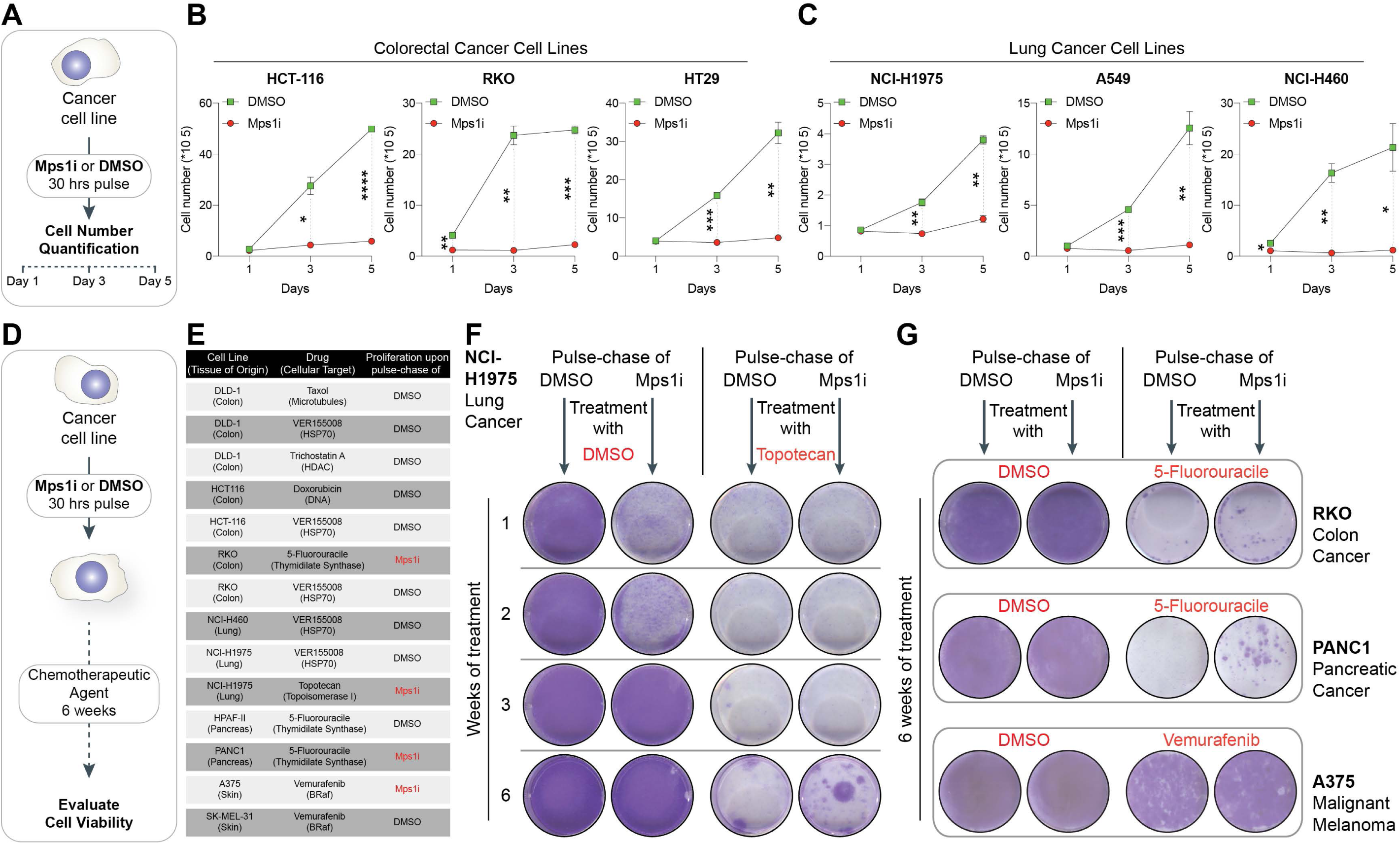
Elevation of chromosome mis-segregation rate facilitates tolerance to chemoresistance. (A) Schematic representation of the experimental set-up utilized to evaluate the effect of chromosome mis-segregation induction on the proliferation of a panel of colorectal and lung cancer cell lines. (B, C) Growth curves of the indicated colorectal (B) and lung (C) cancer cell lines are displayed. Cells were first treated for 30 hours with reversine 0,5μM (red) or DMSO (green). After drug wash-out, cells were plated into multi-well plate and counted 1, 3 and 5 days later. Data represent mean ± SEM from 2 biological replicates; *p<0.05, **p<0.01, ***p<0.001, ****p<0.0001, Student’s t test. (D, E) Schematics of experimental plan (D) and list of cancer cell lines (E) utilized to evaluate viability upon treatment with a chemotherapeutic agent after pulse with reversine 0,5μM or DMSO. In red, conditions under which Mps1i pulse provided a proliferative advantage. Cell viability was evaluated by Crystal Violet staining 6 weeks after continuous treatment with the chemotherapeutic agent. (F-G) Viability assay of NCI-H1975, RKO, PANC1 and A375 pre-treated for 30h with DMSO or reversine and then treated for 6 weeks with 0,1μM Topotecan, 3,5 μM 5-fluoruracile or 1μM vemurafenib. At the end of the treatment, cells were stained with Crystal Violet solution.

Interestingly, the amount of CIN induction required for survival upon chemotherapy was dependent on the tested cell line and the chemotherapeutic agent employed (Supplementary Figure 1A, B – pulse with reversine 500 nM successfully led to the emergence of colonies in NCI-H1975 in the presence of the chemotherapeutic agent, whereas reversine 250 nM pulse was sufficient to achieve similar results in RKO in the presence of the anticancer agent), indicating that an optimal degree of chromosomal instability is required to successfully achieve chemoresistance. Further, we tested whether triggering genome instability by other means not directly related to whole chromosome mis-segregation would also provide a proliferative advantage. For this, we pulsed NCIH1975 with the DNA polymerase inhibitor aphidicolin (Supplementary Figure 1C), the DNA intercalator doxorubicin (Supplementary Figure 1D) or the alkylating agent Cisplatin (Supplementary Figure 1E) before exposure to topotecan. We found that those genome instability-inducing agents did not facilitate the emergence of colonies following treatment with the chemotherapeutic agent, suggesting that karyotypic heterogeneity may be required for chemoresistance in some cell line-drug combinations. Finally, to confirm that these results were not specific to reversine pre-treatment, we pulsed NCI-H1975 with AZ3146 – a different Mps1 inhibitor (Hewitt et al. 2010) - and obtained similar outcomes (Supplementary Figure 1F). Importantly, by using a similar experimental setup and chemically-unrelated Mps1 inhibitors, Lukow et al. also showed that induction of CIN was able to accelerate the generation of resistant cells able to proliferate in the presence of chemotherapeutic agents. Altogether, our results and those by Lukow and co-workers indicate that in spite of the fact that induction of CIN is detrimental for cell proliferation, it might be beneficial for cancer cells under conditions of chemotherapy regimen, thus providing a proliferative advantage.

Interference with the process of chromosome segregation by inhibiting the catalytic function of Mps1 leads to random chromosome gains and losses (Santaguida et al. 2017). While the resulting chromosome imbalances might have detrimental effects on cell physiology (Figure 1A-C), they could also provide phenotypic heterogeneity and might fuel cancer growth, for instance by sculpting the genome through cumulative haploinsufficiency and triplosensitivity (Davoli et al. 2013). We therefore considered that chromosome reshuffling imposed by mitotic errors might expand the karyotypic landscape, thus allowing cancer cells to evolve specific chromosomal assortments that would render them resistant to chemotherapy. To formally test this, we firstly checked the impact of Mps1 inhibitor (Mps1i) treatment on karyotypic heterogeneity and we decided to focus our attention on NCI-H1975, given the fact that chemoresistance is the main cause for therapeutic failure in NSCLC (Chang 2011). Karyotype analysis by single-cell whole-genome sequencing (scWGS) (Figure 2A) showed no major changes over time in terms of heterogeneity score (HS, *i*.*e*. the difference in chromosome copy number between individual cells in a sample (Bakker et al. 2016)) in vehicle-control treated cells compared to the parental line (Figure 2B and Supplementary Figure 2A-E). At the same time, a transient and massive (∼3 fold) increase in HS for all chromosomes was observed right after Mps1i pulse (time 0 in Figure 2C) compared to parental line (average HS: 0.31 in parental, 1.03 in Mps1i wash-out at time 0 - Figure 2C and Supplementary Figure 2A-E). We next examined the karyotypes of NCI-H1975 cells resistant to Topotecan. Analysis of Topotecan-resistant cells either DMSO or Mps1i pulsed (named TRDP - <u>T</u>opotecan <u>R</u>esistant <u>D</u>MSO <u>P</u>ulsed - and TRMP - <u>T</u>opotecan <u>R</u>esistant <u>M</u>psi1 <u>P</u>ulsed -, respectively) as well as parental line revealed that they were all characterized by both segmental and whole-chromosome aneuploidies (Supplementary Figure 2A and 3A-C). Remarkably, a feature stood out whereby the HS of specific chromosomes was lower in resistant cells compared to parental line (Figure 2D and Supplementary Figure 2A and 3A). In particular, chromosomes 4, 6, 15 and 21 in TRMP were those with the lower HS (Figure 2D), indicative of the presence of clonal karyotypes (Supplementary Figure 3A). Interestingly, although the karyotypes of TRDP and TRMP were different (Supplementary Figure 3A), we also found chromosomes 6 and 21 to have a low HS in TRDP, suggesting that the ploidy of those chromosomes might be involved in the acquired chemoresistance. Further, to confirm that emergence of chemoresistance associated with clonal karyotypes was not a unique feature of Topotecan-treated NCI-H1975, we determined the karyotypes of RKO cells resistant to 5FU (Figure 1G) following a DMSO or an Mps1i pulsed and compared them to their respective parental line (Supplementary Figure 4A). Analysis by scWGS showed that gain of chromosome 14 was a major feature of 5-FU resistant RKO pulsed with Mps1i (Supplementary Figure 4A, B), suggesting that chromosome 14 gain might drive chemoresistance to 5FU in RKO cells.

**Figure 2.**
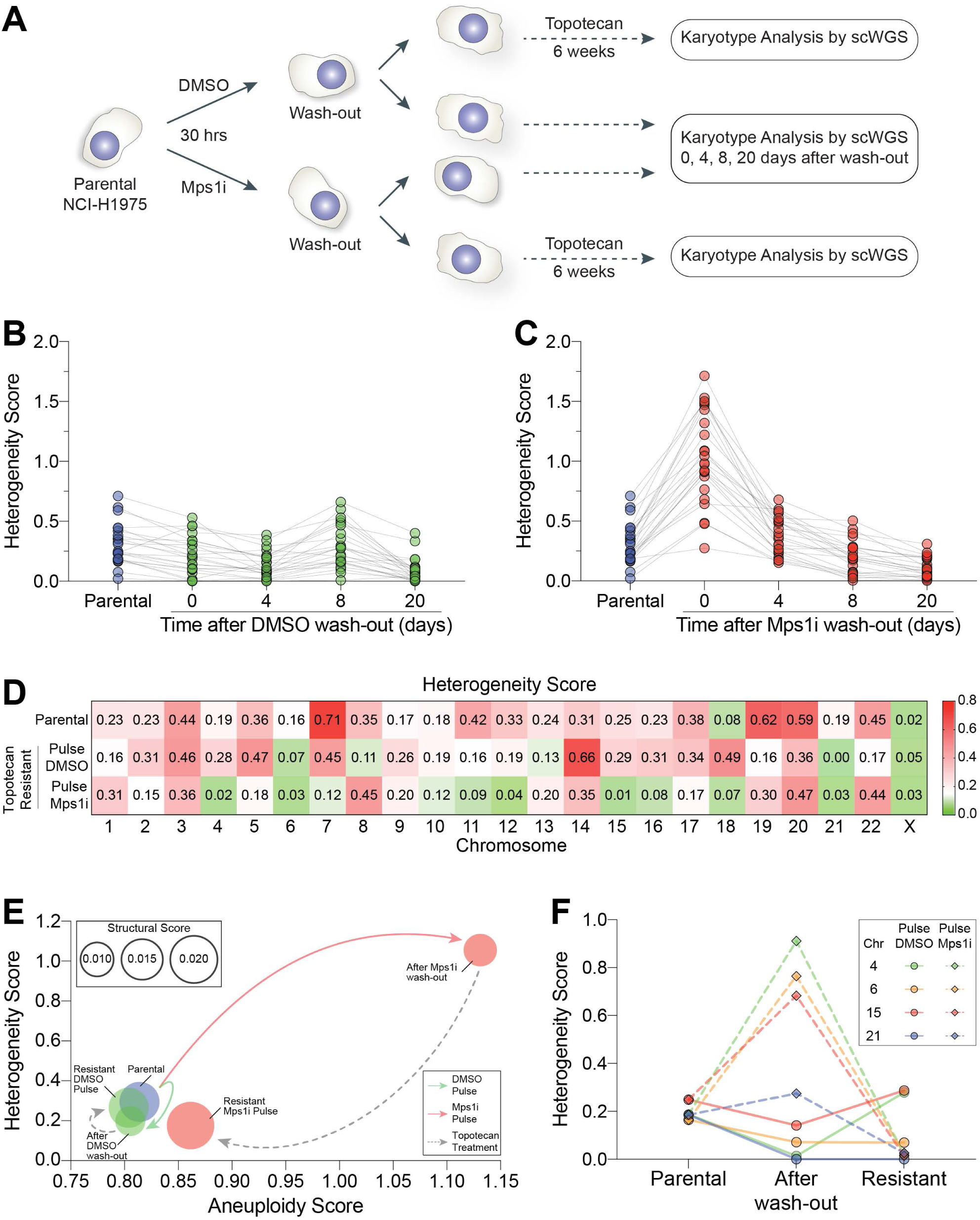
Chemoresistant cells are characterized by the presence of recurrent karyotypes. (A) Workflow for the generation of topotecan-resistant NCI-H1975 cells and their sequencing. NCI-H1975 were treated for 30h with DMSO or reversine. The compound was washed out and cells were either treated with Topotecan for 6 weeks or harvested 0, 4, 8, 20 days after wash-out. In both cases, karyotypes analysis was performed by single-cell whole-genome sequencing (scWGS). (B, C) Heterogeneity scores after DMSO (A) or Mps1i (reversine, B) pulse. Cells were treated as described in (A) and heterogeneity score determined 0, 4, 8, 20 days after wash-out. Parental NCI-H1975 are shown as reference. The lanes are connecting the value of heterogeneity scores for a given chromosome across time points. See also Supplementary Fig. 2. (D) Values of heterogeneity scores for parental NCI-H1975, TRDP and TRMP are shown. See also Supplementary Fig. 2 and 3. (E) Bubble plot showing structural, aneuploidy and heterogeneity scores for NCI-H1975, either before treatment (light blue), or after wash-out from either DMSO and subsequent Topotecan treatment (light green) or Mps1i and subsequent Topotecan treatment (light red). See also Supplementary Fig. 2 and 3. (F) Heterogeneity scores of chromosomes 4, 6, 15 and 21 in parental NCI-H1975 before treatment and right after DMSO or Mps1i wash-out, and in resistant cells.

**Figure 3.**
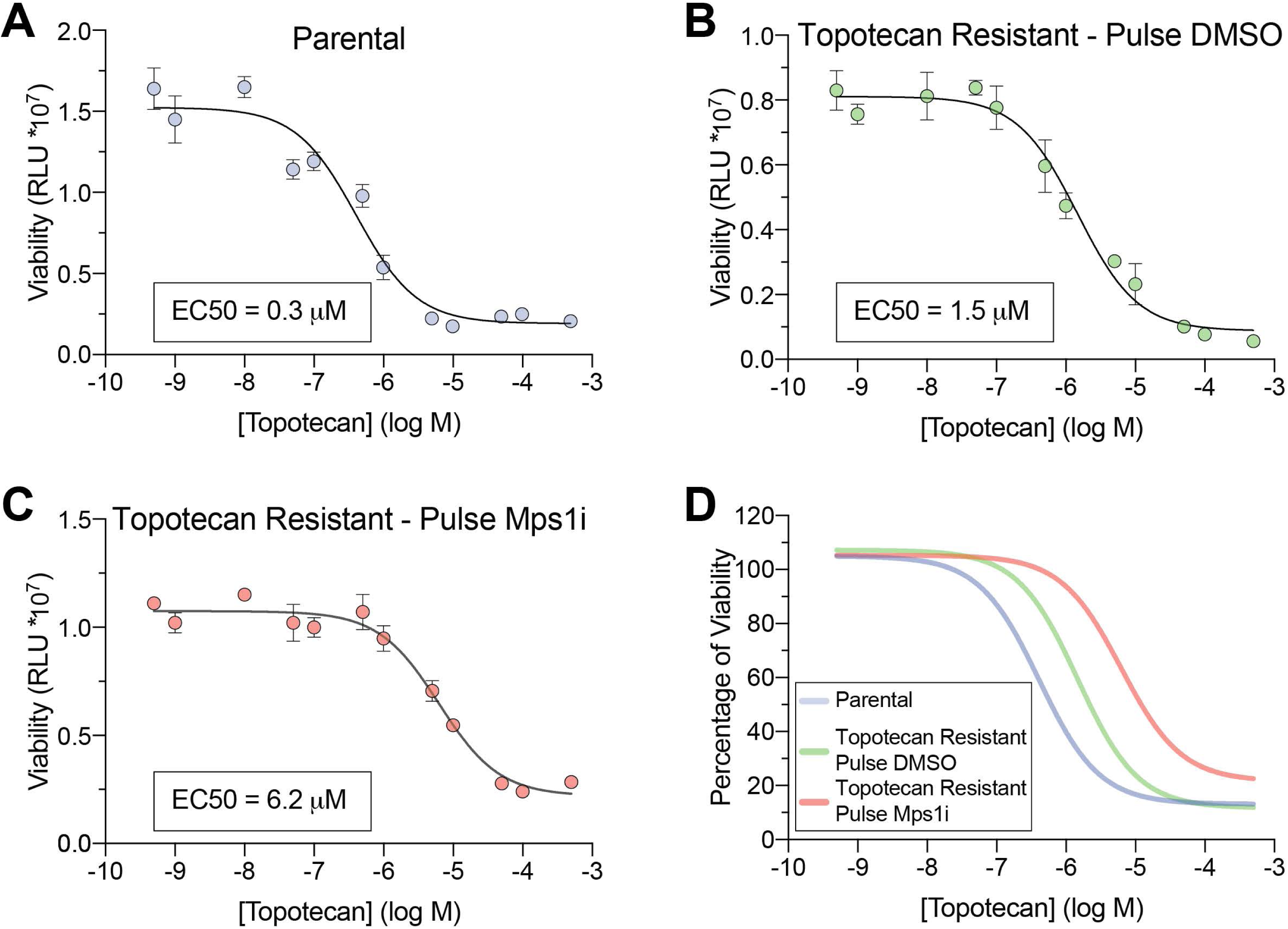
Cancer cells pulsed with Mps1i are more resistant to chemotherapeutic agents. (A-D) Viability assays of Parental (A), TRDP (B), TRMP (C) cells treated with the indicated concentrations of Topotecan for 72 hours and comparison of their profiles (D). The values of calculated half maximal effective concentration (EC50) are shown (A-C).

**Figure 4.**
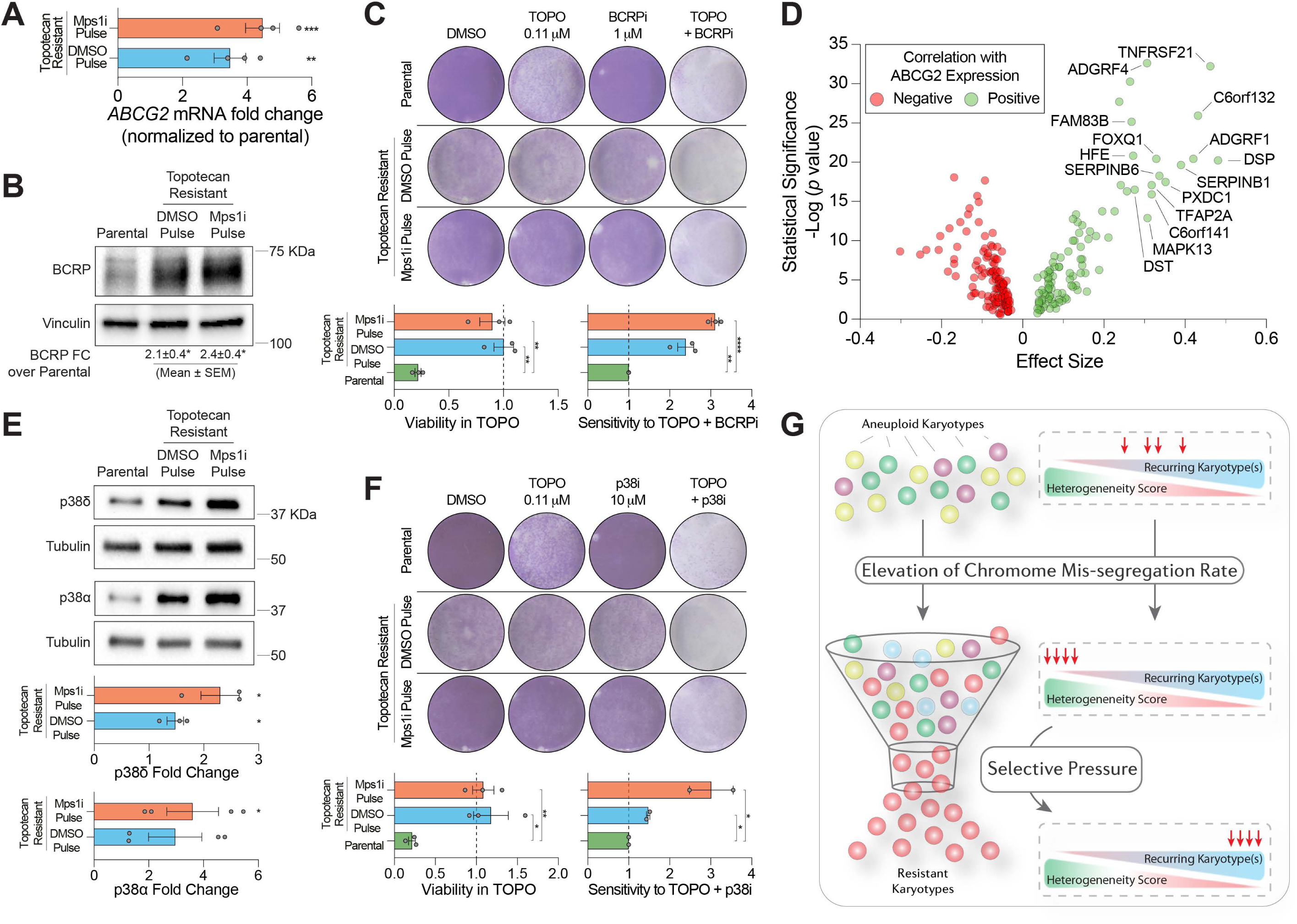
Aneuploidy-induced upregulation of specific proteins drive chemoresistance. (A-B) *ABCG2* mRNA levels and BCRP protein levels (B) were determined in parental NCI-H1975, TRDP and TRMP cells. In (B), Vinculin was used as loading control. (C) Crystal violet staining of parental NCI-H1975, TRDP and TRMP cells after continuous treatment for 5 days with vehicle control (DMSO, first column), Topotecan 0.11 μM (second column), BCRP inhibitor 1 μM (Ko143, third column), Topotecan 0.11 μM and BCRP inhibitor 1 μM (fourth column). Quantification of viability in Topotecan (determined as ratio of crystal violet intensities of Topotecan-treated cells over DMSOtreated) and reacquired sensitivity to Topotecan in presence of BCRPi (measured as ratio of crystal violet intensities of Topotecan-treated cells over Topotecan + BCRPi cotreatment) are quantified on the bottom. (D) A volcano plot showing the co-expression of genes that reside on chromosome 6 with the expression of *ABCG2*. (E) p38delta and p38alpha protein levels were determined in parental NCI-H1975, TRDP and TRMP cells. Tubulin was used as loading control. Quantification is shown on the bottom. (F) Crystal violet staining of parental NCI-H1975, TRDP and TRMP cells after continuous treatment for 5 days with vehicle control (DMSO, first column), Topotecan 0.11 μM (second column), p38 inhibitor 10 μM (SB203580, third column), Topotecan 0.11 μM and p38 inhibitor 10 μM (fourth column). Quantification of viability in Topotecan (determined as ratio of crystal violet intensities of Topotecan-treated cells over DMSO-treated) and reacquired sensitivity to Topotecan in presence of p38i (measured as ratio of crystal violet intensities of Topotecan-treated cells over Topotecan + p38i co-treatment) are quantified on the bottom. (G) A model for how aneuploidy-induced genome instability creates permissive conditions under selective pressure. See text for more details. Data show mean ± SEM; *p<0.05, **p<0.01, ***p<0.001, ****p<0.0001, Student’s t test.

The recurrent patterns of particular chromosomes seen in both NCI-H1975 resistant cell lines could be a consequence of a positive selection - driven by the chemotherapeutic agent - for an underlying karyotype already present in the parental population and/or the result of a process of genome reshaping that drives convergent evolution onto a specific karyotypic state. We reasoned that the former might be a possible scenario for TRDP cells, whereas the combination of the two conditions might reflect what occurred in TRMP. To test this, we analyzed - at the cell population level - aneuploidy, heterogeneity and structural scores, which provide the coordinates of the genomic space into which a cell population can navigate (Figure 2E). Notably, Mps1i treatment imposed a departure from the parental genome, allowing the cell population to reach a different karyotypic state, mainly characterized by increased heterogeneity and aneuploidy scores. This state is inherently unstable (Figure 2C), thus providing the possibility to sample different genomic landscapes and empowering rapid genomic drifts during treatment with the chemotherapeutic agent. Indeed, subsequent exposure to Topotecan allowed the population to select for one particular state, thereby resulting in lower heterogeneity and aneuploidy scores. On the other hand, as expected, pulsing the parental line with DMSO did not significantly impact those parameters and the resultant resistant were as heterogenous and as aneuploids as the starting population (Figure 2E). These observations, made at the level of the entire genome, are mirrored by the analysis of single chromosomes (Figure 2F). In particular, by tracing the evolution of chromosomes with the lower HS in resistant cells (chromosome 4, 6, 15 and 21), we noticed that their HS did not undergo major changes when the parental line was pulsed with DMSO, and a similar trend was observed from DMSO washout to the generation of TRDP (Figure 2F). In striking contrast, HS was increasing following Mps1i treatment, indicative of karyotypic expansion, and then drastically decreased in TRMP, reflecting the convergence on specific karyotypes.

Chromosome mis-segregation has been proposed to be a non-random process (Worrall et al. 2018; Dumont et al. 2019). We therefore considered whether the specific karyotypic remodeling present in TRMP was the result of a biased chromosome mis-segregation pattern caused by Mps1i treatment. Our data suggest that this was not the case, at least for two reasons. First, analysis of scWGS did not show biases towards the missegregation of any particular chromosome when cells were exposed to Mps1i (Figure 2AC, Supplementary Figure 2A, B). Second, exposing cells to Topotecan 6 weeks after Mps1i pulse (rather than right after the pulse and which corresponds to the timeframe required for the formation of resistant clones (Figure 1F)) did not lead to the emergence of colonies (Supplementary Figure 5A). Collectively, our data suggest that CIN might drive chemoresistance leading to the selection of specific karyotypes that become crucial under conditions of selective pressure. Those karyotypes are enabled by the karyotypic heterogeneity induced by Mps1i treatment.

The observation that TRDP and TRMP share some (but not all) recurrent chromosomes prompted us to consider the extent to which the Mps1i pulse provided tangible benefits upon treatment with the chemotherapeutic agent. For this, we calculated the half maximal effective concentrations (EC50) of Topotecan for both TRDP and TRMP as well as for the parental line (Figure 3A-D). We measured an EC50 of 0.3 µM for the parental line (Figure 3A) and EC50 values of 1.5 µM and 6.2 µM for TRDP and TRMP, respectively (Figure 3B, C). The EC50 value of TRDP indicates 5-fold increase over the parental line, whereas the value measured for TRMP shows a 20-fold increase (Figure 3D). These results indicate that reshaping of the genetic landscape induced by Mps1i treatment provided a substantial change in the ability of cells to survive under selective pressure (Figure 1F).

We next focused our attention on the mechanism underlying chemoresistance in TRDP and TRMP. A major determinant of chemoresistance, often seen in cancer cells, is provided by the overexpression of the therapeutic drug target (Holohan et al. 2013). However, when we measured the levels of topoisomerase 1 – the target of Topotecan - we did not find a difference between parental and resistant cell lines (Supplementary Figure 6A). We then considered another well-known mechanism of chemoresistance, which is overexpression of drug efflux pumps (Holohan et al. 2013). Those are transmembrane proteins that include MDR1, MRP1 and BCRP, encoded by *ABCB1, ABCC1* and *ABCG2*, respectively. Efflux pumps have different substrate specificity but share the same mechanism of action, namely an ATP-dependent conformational change mediating the binding to a specific substrate and subsequent extrusion from the cell (Robey et al. 2018). Interestingly, although there was no increase in *ABCB1* and *ABCC1* expression between parental and resistant cell lines (Supplementary Figure 6B; note we were unable to detect any signal for *ABCB1* transcripts in the tested samples), *ABCG2* mRNA levels were increased in both resistant cell lines (Figure 4A), which in turn led to increased protein expression as well (Figure 4B). Importantly, inhibition of BCRP restored sensitivity to Topotecan in resistant cell lines (Figure 4C), indicating that chemoresistance in TRDP and TRMP was mediated by BCRP up-regulation. Intriguingly, the chromosomal region encompassing *ABCG2* – *q* arm of chromosome 4 - was not amplified in TRMP and only in a fraction of TRDP cells (Supplementary Figure 6C). A potential reason for this lack of amplification could be provided by the identity of genes present on this arm of chromosome 4, which have known tumor suppressor functions (*e*.*g*., TET2 [4q24], FAT4 [4q28.1], FBXW7 [4q31.3], FAT1 [4q35.2]). This led us to speculate that because of those genes, resistant cells might have been able to find other ways to upregulate a particular gene of interest, such as *ABCG2*, without amplifying the corresponding chromosomal region, thus avoiding the load of extra copies of tumor suppressors. To identify potential proteins/pathways responsible for *ABCG2* upregulation, we searched for genes whose expression positively correlates with *ABCG2* expression (Figure 4D), by interrogating the Cancer Cell Line Encyclopedia (CCLE) gene expression dataset (Ghandi et al. 2019). To narrow down our search, we imposed two constraints. First, genes had to be located on gained chromosomal regions. Second, those regions had to be in common between TRDP and TRMP. The only chromosome satisfying these constraints was chromosome 6, whose short arm was gained in almost all cells in both conditions (Supplementary Figure 3A and Figure 2D - note that chromosome 21 showed also low HS in both conditions but was not amplified). We therefore focused on the top 15 genes co-expressing with *ABCG2* and residing on chromosome 6 (Figure 4D). Among them, *MAPK13* stood out for its role in Topotecan chemoresistance, and because, together with its homologue *MAPK14*, it is located within the minimal gained region of chromosome 6p in both of the resistant cell lines (Supplementary Figure 6D). *MAPK13* and *MAPK14* encode for the subunits p38delta and p38alpha, respectively, of the stress kinase p38. We found an increase in *MAPK14 mRNA* as well as in p38alpha and p38delta protein levels in resistant cell lines (Supplementary Figure 6E and Figure 4E).

Importantly, inhibition of p38 kinase activity with a chemical inhibitor was able to rescue sensitivity to Topotecan in resistant cells (Figure 4F and Supplementary Figure6F), pointing at p38 as the main driver of chemoresistance in TRDP and TRMP cells. Notably, this dependency on p38 for sustained chemoresistance seemed to be directly linked to BCRP, as p38 inhibition decreased the amount of BCRP localized at the plasma membrane (Supplementary Figure 6G, H).

Taken together, our data suggest a model in which elevation of chromosome missegregation rate in a cell population increases karyotypic heterogeneity (Figure 4G). At this stage, the cell population is highly genomically unstable, a condition that might be unfavorable for cell proliferation (Figure 1A-C), but could provide the ability to widely sample the genomic space and, in the presence of chemotherapeutic agents, eventually find the right karyotypic assortment for chemoresistance (Figure 4G). This might happen through amplification of a chromosomal region encompassing a gene directly involved in chemoresistance, such as the therapeutic drug target or, as in the case presented here with the NCI-H1975 cell line, by overexpressing a gene that upregulates a drug efflux pump. Altogether, our study indicates that, although detrimental for cell proliferation under normal conditions, CIN and the resulting aneuploidy could be exploited by cancer cells to survive under selective pressure (such as chemotherapeutic drugs), which might contribute to aneuploid karyotypes and CIN being a widespread feature of advanced tumors.

Interestingly, our current results in cell lines mimic our previous results in patient-derived xenografts (PDX), where we found elevated levels of CIN to be associated with chemoresistance (Ben-David et al. 2017). PDXs, however, do not lend themselves easily to functional manipulations. Using cell lines, we were now able to follow the karyotypic evolution of cancer cells exposed to chemotherapeutic agents, and demonstrate that the proliferation advantage induced by CIN under conditions of selective pressure is mediated by the selection of an optimal karyotype. For Topotecan resistance in NCI-H1975 cells, we provide a molecular link between the selected karyotype and the acquired chemoresistance.

We conclude that aneuploidy and CIN are strong promoters of phenotypic variation. Current therapeutic approaches focus on drugs that increase CIN (Pauer et al. 2004; Mason et al. 2017; Wang et al. 2019). While these approaches can be valuable, our results demonstrate that developing approaches to decrease CIN (*e*.*g*., (Orr et al. 2016)) are equally important. More research is required to determine which patients may benefit from each of these opposite strategies, which will be instrumental for overcoming chemoresistance.

## Acknowledgements

We are grateful to Devon Lukow and Jason Sheltzer (CSHL) for sharing data prior to publication. We thank Stephen Taylor (University of Manchester) and members of the Santaguida lab for constructive discussions throughout the project. Work in the Santaguida lab is supported by grants from the Italian Association for Cancer Research (MFAG 2018 - ID. 21665 project), Ricerca Finalizzata (GR-2018-12367077), Fondazione Cariplo, the Rita-Levi Montalcini program from MIUR and the Italian Ministry of Health with Ricerca Corrente and 5×1000 funds.

## Supplementary Figure Legends

**Supplementary Figure 1**. (A, B) Viability assay of NCI-H1975 (A) and RKO (B) cell lines in presence of Topotecan and 5FU, respectively, after pulse with DMSO or reversine. Cells were treated for 30h with DMSO or reversine at different concentrations (0.125, 0,25, 0,5 μM). The compounds were then washed out and Topotecan 0.11 μM (to NCI-H1975) or 5fluoruracile 3.5 μM (to RKO) were added. Cell viability was evaluated by crystal violet staining 4, 12, 24 and 36 days later. (C-F) NCI-H1975 were treated for 30h with DMSO or: aphidicolin 0,4μM, 1μM (C), doxorubicin 0.1μM, 0.2μM (D), cisplatin 1μM, 2μM (E), and AZ3146 1μM, 2μM (F). Then, the drugs were removed and Topotecan 0,11μM was added for 36 days. At the end of the treatment, cells were stained with Crystal Violet solution.

**Supplementary Figure 2**. (A-E) Genome-wide copy number profiles of populations of parental NCI-H1975 (A), or after pulse of DMSO or Mpsi1 (B-E). Cells were treated as described in Figure 2A and harvested 0 (B), 4 (C), 8 (D), 20 (E) days after wash-out. Single cells are represented in rows and chromosomes plotted as columns. Copy number states are indicated in colors (see Legend on the bottom).

**Supplementary Figure 3**. (A) Genome-wide copy number profiles of populations of TRDP and TRMP cells. Single cells are represented in rows and chromosomes plotted as columns. Copy number states are indicated in colors (see Legend on the bottom). (B, C) Aneuploidy (B) and structural scores (C) of parental NCI-H1975, TRDP and TRMP cells.

**Supplementary Figure 4**. (A) Genome-wide copy number profiles of populations of RKO 30 hours after DMSO or Mps1 pulse, or 5-FU resistant previously pulsed with DMSO or Mps1i. Single cells are represented in rows and chromosomes plotted as columns. Copy number states are indicated in colors (see Legend on the bottom). (B) Cumulative plots of 5-FU resistant RKO cells previously pulsed with Mps1i showing (on the left) or not (on the right) recurrence of chromosome 14.

**Supplementary Figure 5**. (A) NCI-H1975 were treated with DMSO or reversine 0,5μM for 30h. After wash out, the cells were cultured for 6 weeks. At this point, topotecan was added for 36 days and cell viability evaluated by crystal violet staining every 12 days. Under these conditions, no resistant colonies were found.

**Supplementary Figure 6**. (A) Western Blot analysis of Topoisomerase 1 protein levels in parental, DMSO and Mps1i Pulse resistant cells. Tubulin was used as loading control. (B) *ABCC1* mRNA levels were determined in parental NCI-H1975, TRDP and TRMP cells. (C, D) Chromosomal location of *ABCG2* (4q22.1; C), *MAPK13* and *MAPK14* (6p21.31; D) in relation to copy state number of TRDP and TRMP. Copy number states are indicated in colors (see Legend on the bottom). Shown are chromosomes 4 and 6 and are the same sequencing results displayed in Supplementary Figure 3A. (E) *MAPK14* and *MAPK13* mRNA levels were determined in parental NCI-H1975, TRDP and TRMP cells. (F) Crystal violet staining of parental NCI-H1975, TRDP and TRMP cells after continuous treatment for 5 days with vehicle control (DMSO, first column), Topotecan 0.11 μM (second column), p38 inhibitor 1 μM (BIRB796, third column), Topotecan 0.11 μM and p38 inhibitor 1 μM (fourth column). Treatment with p38i rescued sensitivity to Topotecan in TRDP and TRMP cells (compare second and fourth column). (G) Western Blot analysis of BCRP in parental, TRDP and TRMP cells in absence (first three lanes) or presence (last three lanes) of p38i SB203580 after purification of plasma membrane-associated proteins. EGFR was used as control for plasma membrane purification. (H) Quantification of BCRP plasma membrane levels normalized to parental NCI-H1975. Numbers show reduction of BCRP levels in presence of p38i. Data show mean ± SEM; *p<0.05, ***p<0.001, Student’s t test.

## Materials and Methods

### Cell culture condition and reagents

All cell lines were tested free of mycoplasma contamination using Myco Alert (Lonza, Walkersville, MD, USA) according to the manufacturer’s protocol. All cells were maintained in a humidified environment at 37□°C with 5% CO_2_ and cultured in standard medium conditions.

### Drug treatments

Reversine was obtained from Cayman Chemical and used at a working concentration of 0,25 μM or 0,5 μM; Topotecan (working concentration 0,11 μM), Trichostatin A (working concentration 0,5 μM), VER 155008 A (working concentration 5 μM), Taxol (working concentration 0,01 μM) were purchased from Tocris; Vemurafenib (working concentration 1 μM) was purchased by Selleckchem; Doxorubicin (working concentration 0,2 μM), Aphidicolin (working concentration 0,4 μM) and AZ3146 (working concentration 1 or 2 μM) were purchased from Cayman Chemical; SB203580 (working concentration 10 μM) was purchased from Tocris; Cisplatin (working concentration 1, 2 or 3,3 μM) and 5-Fluoruracile (working concentration 3,5 or 9 μM) were obtained from the hospital pharmacy at the European Institute of Oncology (Milan, Italy).

### Cell proliferation assay

HCT-116, RKO, HT29, NCI-H1975, A549, NCI-H460 cells at 50% of confluence, were treated with DMSO or Reversine (0,5μM) for 30h and then the drugs were washed out. After 12h, cells were plated in a 6 well plate and were counted 1, 3 and 5 days after plating using the Bürker counting chamber (Blaubrand, Germany). The experiment was performed in two replicates.

### RNA extraction, RT–PCR and qPCR

RNA was extracted from cells using RNeasy Plus Mini Kit (QIAGEN), according to manufacturer’s protocol. 500 ng of RNA from each sample was reverse-transcribed using OneScript^®^ Plus cDNA Synthesis Kit (abm) according to the manufacturer’s instructions. mRNA expression was performed by real-time quantitative PCR reactions using Fast SYBR™ Green reaction mix (Thermo Fisher Scientific) and achieved on an Applied Biosystems 7500 Fast Real-time PCR system. The relative expression level was calculated with the 2^[DDCt]^ method and expressed as a ‘‘fold change’’: normalization of data was performed on house-keeping gene (*GAPDH*) expression and compared to the Parental cells as a control. Primers used for profiling the mRNA expression levels of genes are as follows: *MAPK14* Fwd: 5-TGCACATGCCTACTTTGCTC-3; Rev: 5AGGTCAGGCTTTTCCACTCA-3; *MAPK13* Fwd: 5GGGATGGAGTTCAGTGAGGA-3; Rev: 5-GTCCTCATTCACAGCCAGGT-3; *ABCB1* Fwd: 5GCCTGGCAGCTGGAAGACAAATAC-3; Rev: 5ATGGCCAAAATCACAAGGGTTAGC-3; *ABCC1* Fwd: 5TGTGTGGGCAACTGCATCG-3; Rev: 5GTTGGTTTCCATTTCAGATGACATCCG-3; *ABCG2* Fwd: 5CCGCGACAGCTTCCAATGACCT-3; Rev: 5-GCCGAAGAGCTGCTGAGAACTGTA-3.

### Protein detection by Western blots

For protein analyses, cells were lysed in RIPA 1x lysis buffer (RIPA buffer 10x; CellSignalingTechnology) with the addition of protease inhibitor cocktail (Millipore), phosphatase inhibitor cocktail (Roche) and then sonicated. Protein lysates were centrifuged at maximum speed for 10 min and resolved on 15% SDS-PAGE gels. The following primary antibodies were used: anti- p38⍰ MAPK (#9218; CellSignalingTechnology, 1:1000), anti- p38δ MAPK (#2308; CellSignalingTechnology; 1:1000), anti-Phospho-p38 MAPK (Thr180/Tyr182) (#9211; CellSignalingTechnology; 1:1000), anti-Topoisomerase 1 (#ab2454311; abcam; 1:1000), anti- ABCG2 (D5V2K) XP Rabbit mAb (#42078; CellSignalingTechnology; 1:1000), anti-GAPDH (#2118; CellSignalingTechnology; 1:1000), anti-Vinculin (#V9131; Sigma-Aldrich; 1:1000), antiTubulin (#T9026; Sigma-Aldrich; 1:1000), anti-EGFR (1:5000; rabbit polyclonal antibody, gift of Pier Paolo Di Fiore).

### Crystal Violet Assay

For Crystal Violet staining, cells were washed twice with ice-cold PBS and fixed with icecold 4% PFA for 15 min. Afterward, 1% crystal violet solution (Sigma V5265) was added to the plates and incubated at room temperature for 15 min. Plates were then washed with distilled H_2_O, until the unbound crystal violet was removed and plates were dried at room temperature. To quantify Crystal Violet intensities, plates were exposed to 10% acetic acid at room temperature for 30 min, on a shaker. Then, the absorbance of the solubilized crystal violet derived from each single well was measured at a wavelength of 600 nm.

### EC50 assay

Cells were plated in 96-well plates (white flat bottom; ThermoFisher Scientific), in triplicate in 50 μl of medium. The following day, cells were treated with drugs dissolved in 50 μl of medium. After 72h of treatment, 100 μl of CellTiter-Glo reagent (Promega; Madison, WI) was added to each well; the plates were incubated at room temperature for 10 minutes and the luminescence signal was measured with a GloMax® microplate reader (Promega; Madison, WI).

### Plasma Membrane purification

Cells were plated in 150mm plates, treated and cultured until 85-90% confluent. Plasma Membrane purification was performed using Pierce™ Cell Surface Protein Biotinylation and Isolation Kit following manufacturer’s instructions for adherent cells; in the lysis step, protease inhibitor cocktail (Millipore) and phosphatase inhibitor cocktail (Roche) were added. Eluted proteins were quantified using Pierce™ BCA Protein Assays. Then, sample buffer was added to eluates and samples were analyzed by Western Blot.

### Sample processing for single-cell sequencing

For single nuclei isolation, cell pellets were resuspended in lysis buffer (1M tris-HCl pH7.4, 5M NaCl, 1M CaCl2, 1M MgCl2, 7.5% BSA, 10% NP-40, ultra-pure water, 10 mg/ml Hoechst 33358, 2mg/ml propidium iodide) and kept on ice in the dark for 15 min to facilitate lysis. G1 single nuclei, as assessed by PI and Hoechst staining were sorted into 96 wells plates on a BD FacsJAZZ cell sorter (BD Biosciences) and stored in -80C until further analysis. For single cell libraries preparation, single nuclei were lysed and DNA was barcoded, followed by automated library preparation (Bravo Automated Liquid Handling Platform, Agilent Technologies) as described previously (van den Bos et al. 2018). Single cell libraries were pooled and analyzed on an Illumina Hiseq2500 sequencer.

### Data analysis single-cell sequencing

Sequencing was performed using a NextSeq 500 machine (Illumina; up to 77 cycles; single end). The generated data were subsequently demultiplexed using sample-specific barcodes and changed into fastq files using bcl2fastq (Illumina; version 1.8.4). Reads were afterwards aligned to the human reference genome (GRCh38/hg38) using Bowtie2 (version 2.2.4; (Langmead and Salzberg 2012)). Duplicate reads were marked with BamUtil (version 1.0.3; (Jun et al. 2015)). The aligned read data (bam files) were analyzed with AneuFinder (Version 1.14.0; (Bakker et al. 2016)). Following GC correction and blacklisting of artefact-prone regions (extreme low or high coverage in control samples), libraries were analyzed using the dnacopy and edivisive copy number calling algorithms with variable width bins (binsize: 1 Mb; stepsize: 500 kb) and breakpoint refinement (R= 20, confint = 0.95; other settings as default). Results were afterwards curated by requiring a minimum concordance of 95% between the results of the two algortithms. Libraries with less than five reads per bin per chromosome copy (∼ 30,000 reads for a diploid genome) were discarded. Samples with a near tetraploid DNA content were analyzed with the developer version of AneuFinder (Version 1.7.4; from GitHub). Depending on the sample, the min.ground.ploidy parameter was set to either 3 or 3.5 and the max.ground.ploidy parameter to 4.5, 5.0 or 5.5. The minimum and maximum ground ploidy values were determined with the results that were previously obtained with the standard (Bioconductor) version of AneuFinder. Results were subsequently curated as described above (except using a minimum concordance of 90%). Aneuploidy and heterogeneity scores were calculated as described in the AneuFinder paper (Bakker et al. 2016).

### Co-expression analysis of *ABCG2* with genes residing on chromosome 6

mRNA expression levels were obtained from the CCLE gene expression data set (19q4 DepMap release; CCLE_mutations.csv) (Ghandi et al. 2019). A gene expression linear association analysis was performed in the DepMap portal (https://depmap.org/portal/), using ABCG2 expression as the dependent variable and the dataset as the independent variable. The analysis regresses a dependent variable on an independent variable and reports a moderated regression coefficient along with its p-value and q-value.

