## Supplementary Figures for "Aneuploidy-driven genome instability triggers resistance to chemotherapy"

Ippolito_Supplementary Fig1

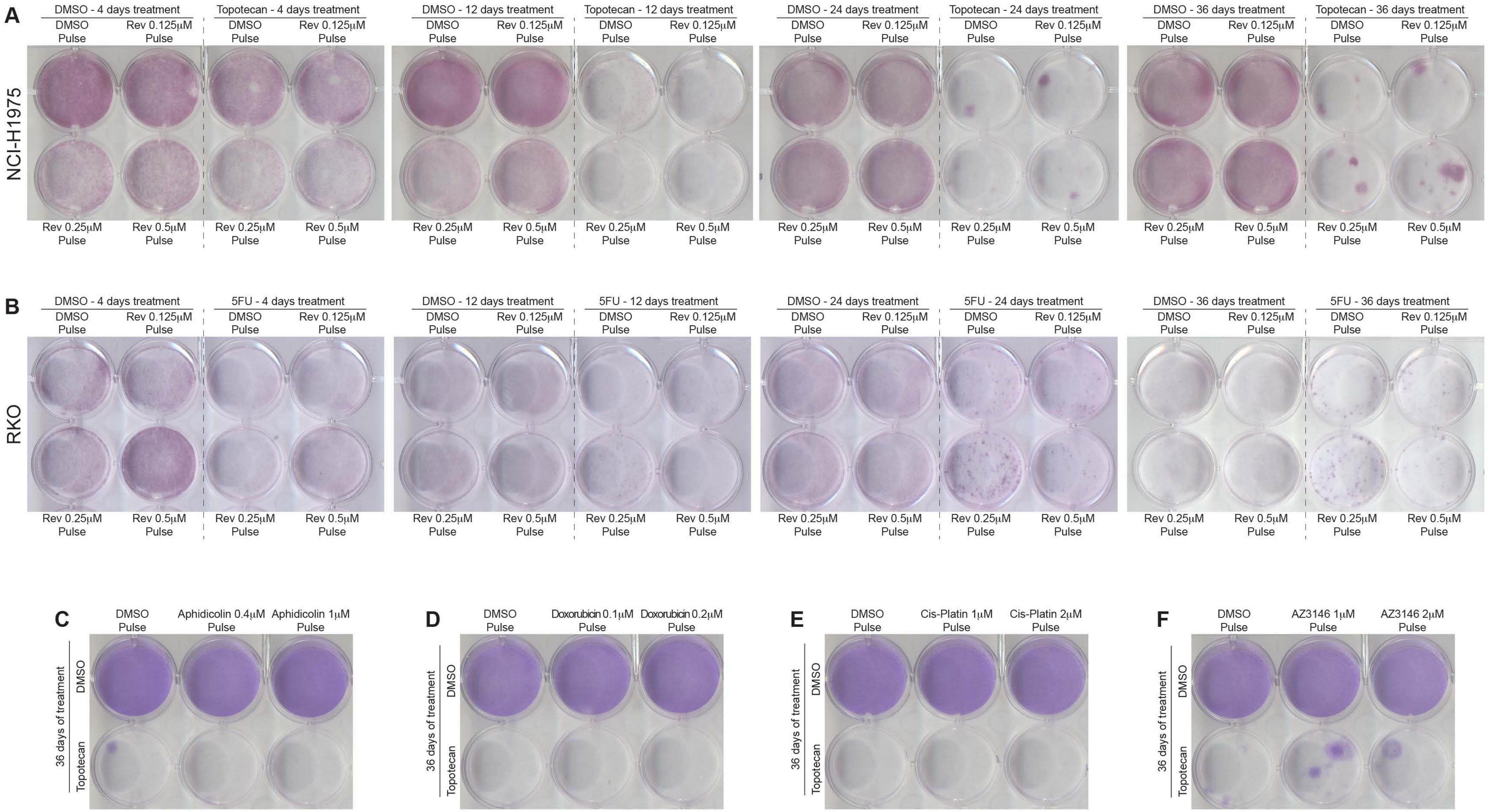

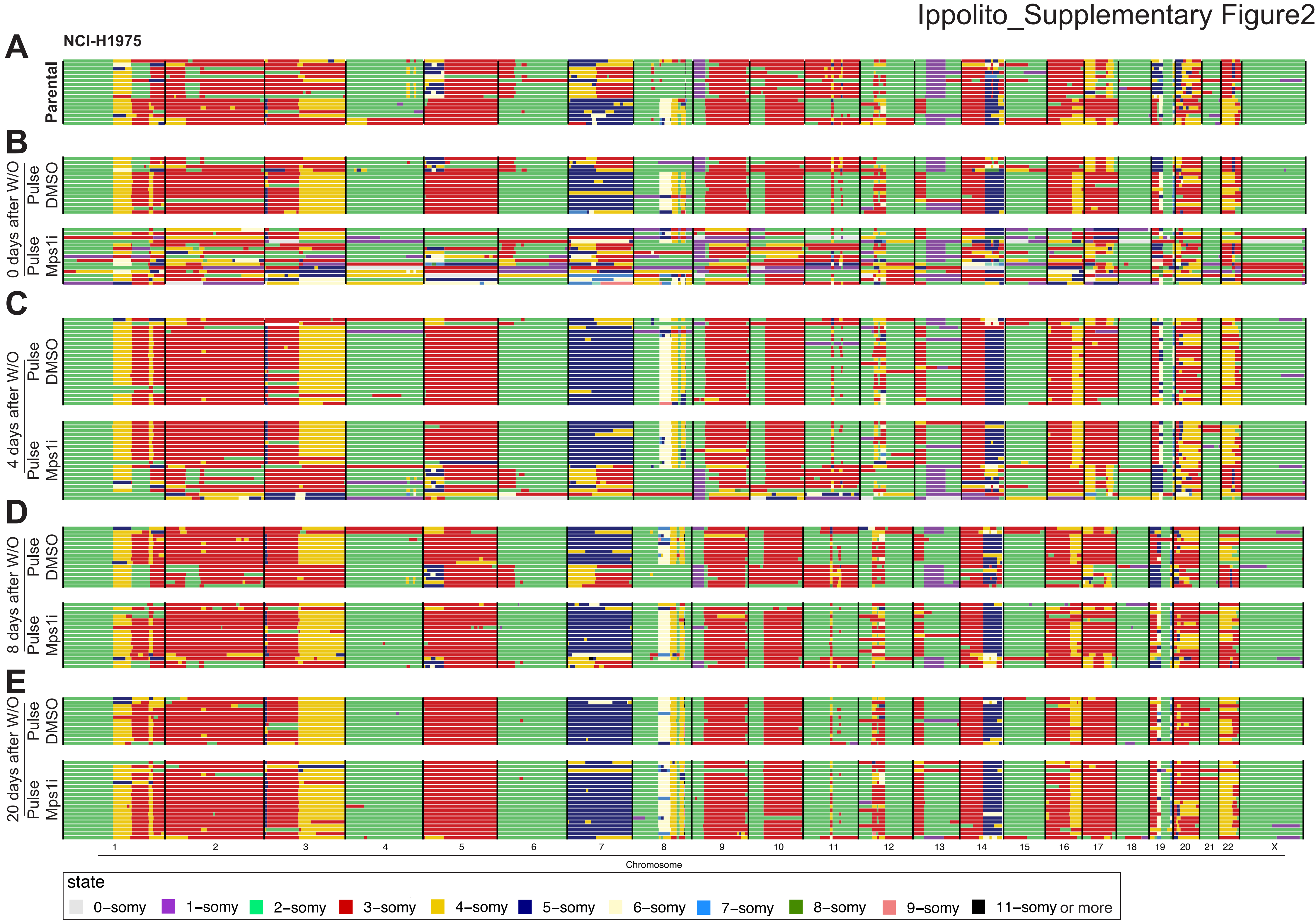

Ippolito_Supplementary Figure2
A
NCI-H1975
Parental
B
0 days after W/O
Pulse DMSO
Pulse Mps1i
C
4 days after W/O
Pulse DMSO
Pulse Mps1i
D
8 days after W/O
Pulse DMSO
Pulse Mps1i
E
20 days after W/O
Pulse DMSO
Pulse Mps1i
1 2 3 4 5 6 7 8 9 10 11 12 13 14 15 16 17 18 19 20 21 22 X
Chromosome
state
0−somy
1−somy
2−somy
3−somy
4−somy
5−somy
6−somy
7−somy
8−somy
9−somy
11−somy or more

Ippolito_Supplementary Fig3

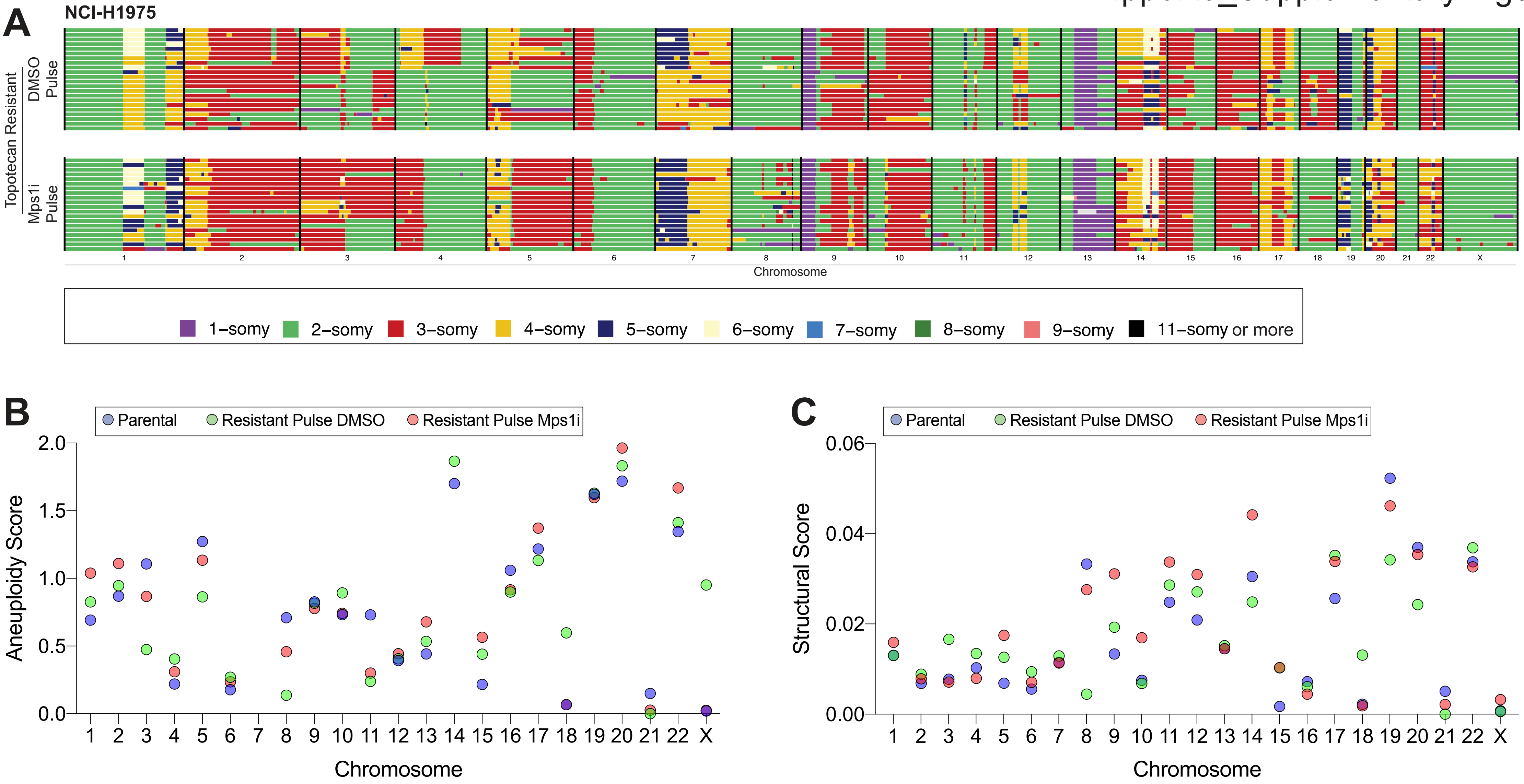

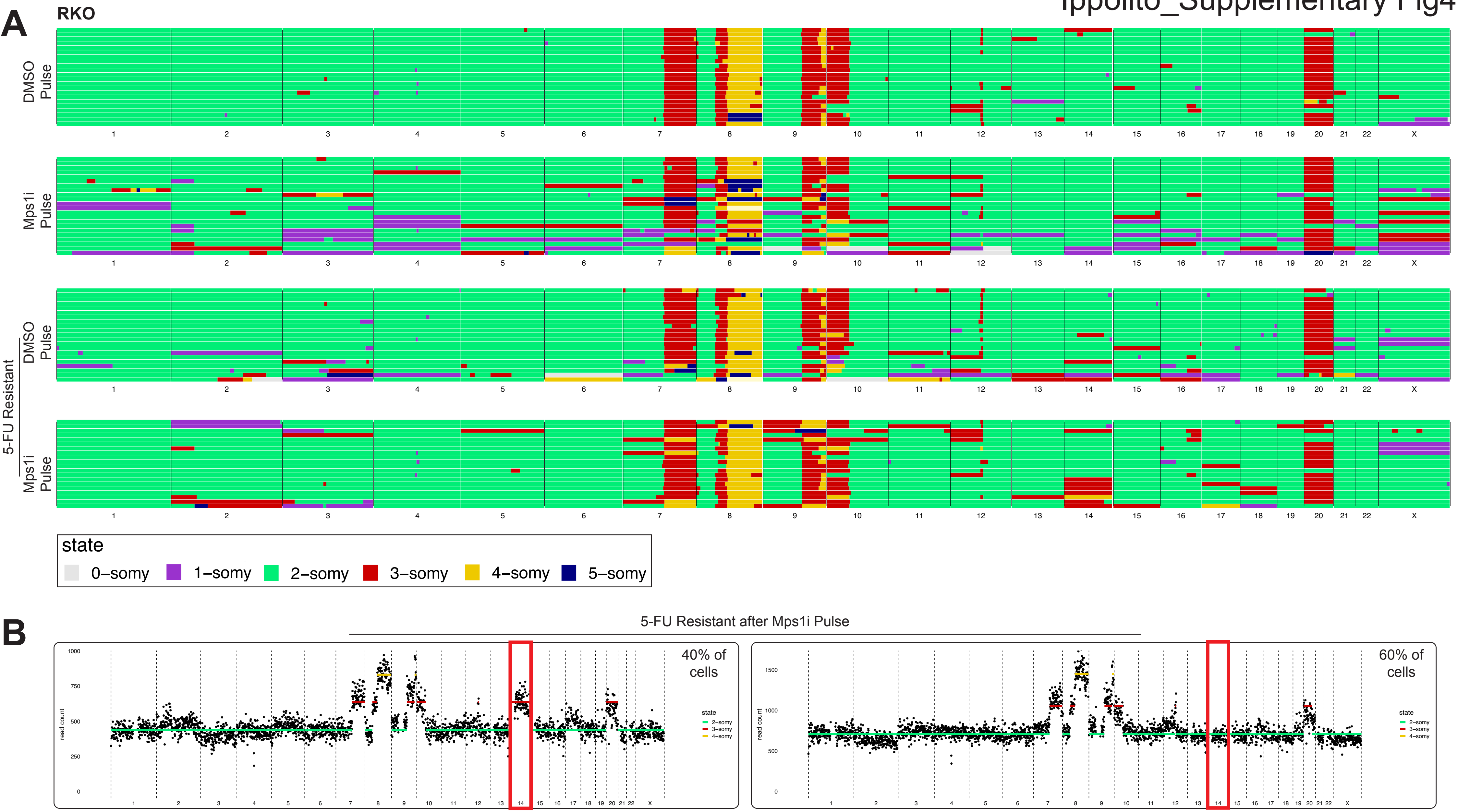
A
RKO
DMSO Pulse
Mps1i Pulse
5-FU Resistant
DMSO Pulse
Mps1i Pulse
1 2 3 4 5 6 7 8 9 10 11 12 13 14 15 16 17 18 19 20 21 22 X
state
0–somy
1–somy
2–somy
3–somy
4–somy
5–somy
B
5-FU Resistant after Mps1i Pulse
40% of cells
60% of cells
read count
state
2–somy
3–somy
4–somy

Ippolito_Supplementary Fig5

A

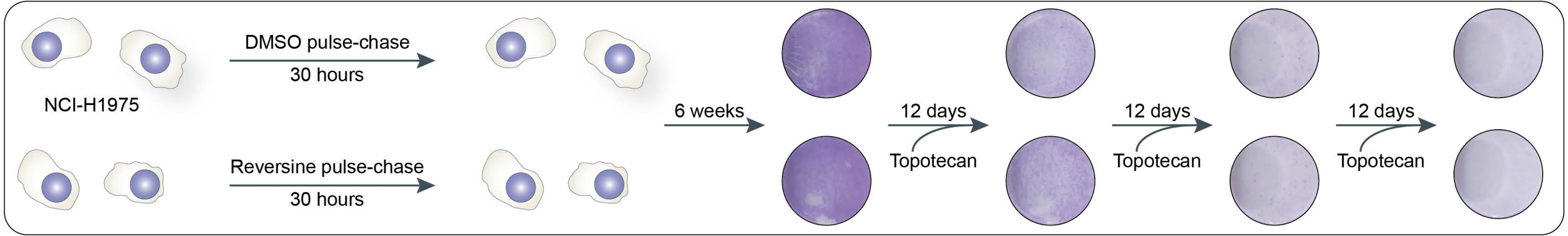

Ippolito_Supplementary Fig6

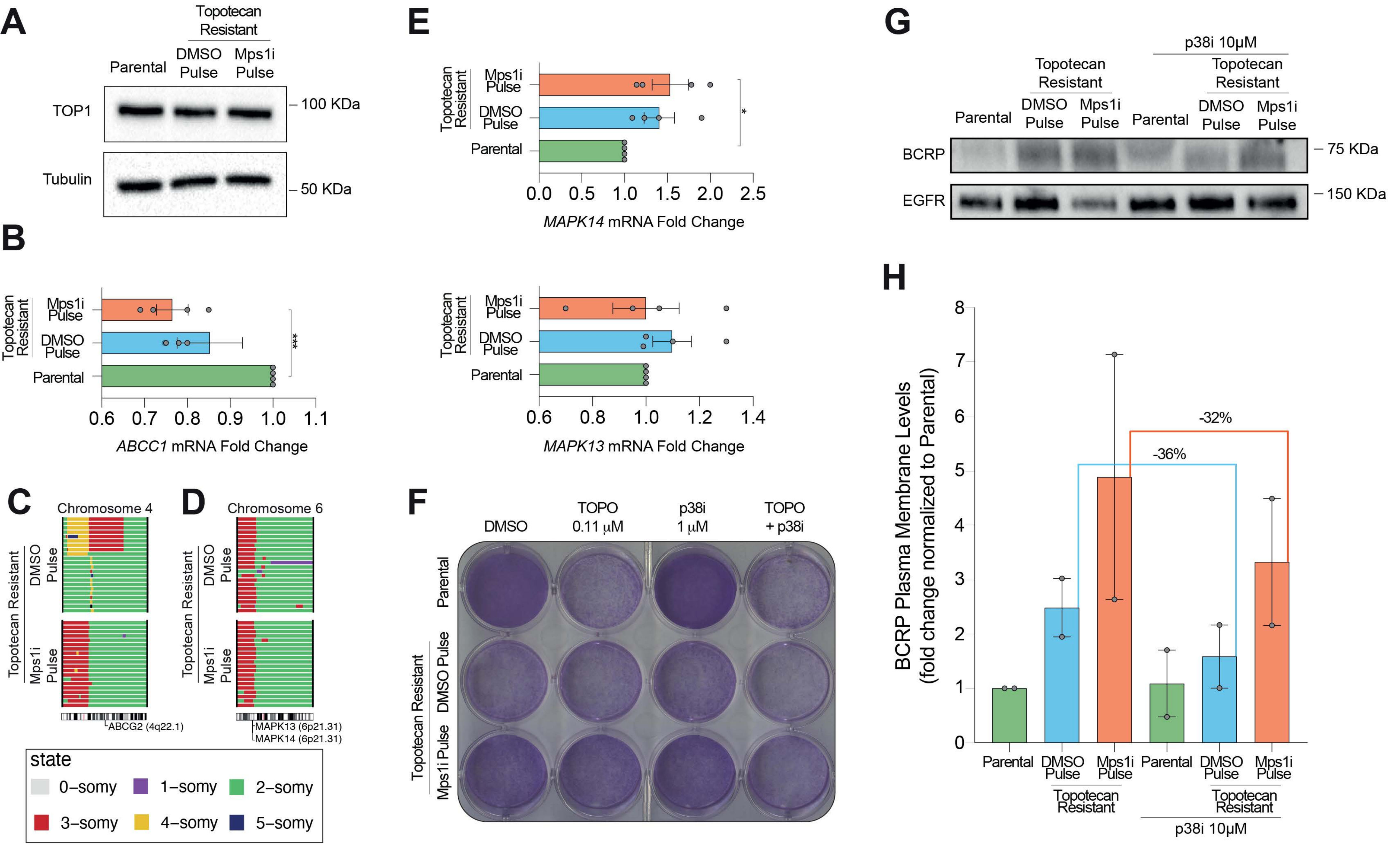
